# Perturbation of nuclear proteins by optogenetic trapping in *Drosophila* ovary

**DOI:** 10.1101/2025.10.07.680910

**Authors:** Bing Liu, Aurelien Guillou, Sijia Zhou, Hao Li, Yaru Kong, Jiaying Liu, Denis Jullien, Thomas Mangeat, Luisa Di Stefano, Xiaobo Wang

## Abstract

The eukaryotic nucleus hosts transcription factors and other numerous nuclear proteins essential for genome organization and function. Optogenetic tools have been applied to match nuclear protein dynamics but remain largely confined to transcription factors, leaving most nuclear proteins inaccessible for precise manipulation. Here, we engineered an optogenetic tool, NLS-LARIAT, by fusing a nuclear localization sequence to the LARIAT (light-activated reversible inhibition by assembled trap), enabling light-inducible clustering of diverse GFP-tagged nuclear proteins in *Drosophila* ovarian cells. NLS-LARIAT–mediated clustering of transcriptional and epigenetic regulators, such as Slbo and LSD1, phenocopied genetic inhibition while revealing rapid and localized roles of Slbo during border cell migration. Beyond regulatory factors, NLS-LARIAT can also be used to manipulate nuclear structures including the nuclear envelope and chromatin. Thus, NLS-LARIAT provides a versatile and broadly applicable optogenetic platform for spatiotemporal control of nuclear proteins, offering new opportunities to dissect nuclear organization and function across model systems.

## Introduction

Diverse cellular and tissue processes are tightly regulated by key intracellular factors and signaling cascades that span the plasma membrane to the nucleus. Proper tissue structure and homeostasis depend on the precise spatiotemporal regulation of these factors and pathways, whereas their dysregulation drives diseases, including cancer (*1–3*). Within the nucleus, gene expression is controlled not only by transcriptional factors but also by nuclear structures such as chromatin state, nuclear envelopes, and nuclear F-actin networks, which couples to both coding and non-coding RNA transcription (*3, 4*). These regulatory activities are highly dynamic and context dependent, varying across developmental stages and tissues (*5*). Traditional genetic methods often have limitations in studying the dynamic spatiotemporal roles of target proteins (*5*). As an alternative tool, optogenetic approaches can precisely manipulate protein functions with higher temporal and spatial resolution (*5*).

Similar to cytoplasmic proteins, nuclear proteins are also highly dynamic, and their spatiotemporal control is pivotal for many cellular processes (*4*). Currently, there are two major optogenetic strategies for photo-manipulating transcription factors. The first strategy is to use light to switch the nuclear entry and exit of a specific transcription factor (*6–15*), and this strategy has also been applied to photo-manipulate a chromatin regulator Sin3A (*14*). The second strategy is to use light to combine the DNA-binding domain and transactivation domain of a specific transcription factor, thereby achieving transcriptional activation/repression of its targets (*16, 17*). Both strategies are limited to a specific transcription factor, as previously reviewed (*5*). Different optogenetic constructs are required for different transcription factors, while common optogenetic tools suitable for different targets are still lacking. Despite significant progress, both strategies fail to control other nuclear factors and structures, such as histones, nuclear envelopes, nuclear lamina, and nuclear bodies, which contribute not only to gene transcription but also to other essential nuclear functions. Thus, there is an urgent need to establish a common optogenetic tool applicable to various nuclear factors.

To overcome the target-specific limitation, we previously established GFP-targeted “light activated reversible inhibition by assembled trap” (LARIAT), a versatile photo-inactivation method, in the *Drosophila in vivo* system (*18, 19*). The working mechanism of GFP-LARIAT is that upon blue light, CRY2 (fused to anti-GFP nanobody) heterodimerization with CIB1 (fused to multimeric protein (MP)) robustly sequesters various GFP-tagged proteins (*20*). This rapidly inactivates targets in the cytoplasm. Using GFP-LARIAT, we and other teams succeeded in the photo-inhibition of myosin contractility and cell-matrix/cell-cell adhesions (*18*), ROCK and phosphatase (*19*), the apical polarity protein aPKC (*21*), and cytoskeleton microtubules (*22*) in different types of *Drosophila* epithelial cells *in vivo*. However, in the original GFP-LARIAT construct, CIB1-MP is unable to enter the nucleus and is not expected to interact with nuclear Cry2-VHH(GFP), thus failing to efficiently photo-inhibit GFP-tagged nuclear targets (*18*).

To overcome this limitation and develop a broad optogenetic tool applicable to nuclear proteins, here we introduced an NLS (nuclear localization sequence) into the CIB1-MP partner, generating NLS-LARIAT for targeting nuclear factors *in vivo*. NLS-LARIAT enables robust clustering of various GFP-tagged nuclear factors in different cell types of the *Drosophila* egg chamber. The clustering-trap−mediated photo-inhibition of GFP-tagged transcriptional and epigenetic factors phenocopies the genetic inhibitory effects, while local and gradual photo-inhibitions provide insights into their spatiotemporal roles in controlling border cell migration. Moreover, NLS-LARIAT extends beyond transcription/epigenetics, which allows spatiotemporal control of other nuclear structures, such as the nuclear envelope and chromatin.

## Results

### NLS-LARIAT clusters various nuclear factors in different *Drosophila* cells

In our previous GFP-LARIAT construct, CIB1-MP cannot enter the nucleus (*18*); consistently, CIB1-MP tagged with mCherry was mainly located in the cytoplasm (fig. S1A). Since the success of LARIAT is highly dependent on the equal amount of CIB1-MP and CRY-VHH(GFP) expression levels (*20, 23*), we used the 2A linker system (*24*) to drive the expression of both CIB1-MP and CRY-VHH(GFP) in previous GFP-LARIAT. To produce GFP-LARIAT suitable for nuclear proteins, we inserted a nuclear localization signal (NLS) into 6 different regions of this GFP-LARIAT construct (schematic in fig. S1B). When using 3 constructs (#1, #2, #5), we still detected mCherry tagged CIB1-MP in the cytoplasm; whereas for the other 3 constructs (#3, #4, #6), mCherry tagged CIB1-MP was strongly located within the nucleus (fig. S1C). We then compared the effects of GFP clustering by these 6 constructs vs. the previous GFP-LARIAT in *Drosophila* S2 cells. For constructs (#1, #2, #5), 4-hour visible light illumination mildly clustered nls-GFP and we still detected prominent non-clustered nuclear GFP signals, similar to the effect of previous GFP-LARIAT; whereas for constructs (#3, #4, #6), 4-hour light illumination strongly clustered nls-GFP (fig. S1C). Given the strong clustering trap ability of nuclear GFP signals, we named these 3 constructs (#3, #4, #6) as the new optogenetic tool NLS-LARIAT (Schematic in Fig. 1A) to distinguish it from our previous GFP-LARIAT (which we renamed Cyto-LARIAT). Next, we compared the dynamic response of construct#3, #4, #6 to a pulse of blue light illumination (see Methods). These 3 constructs exhibited different clustering effects after single pulse light illumination: 1) Construct #3 had a slow light response and could not recover in darkness; 2) Construct #4 had a moderate light response and weak recovery in darkness; 3) Construct #6 had a rapid light response and partial recovery in darkness (fig. S2). Considering that construct #6 exhibited the fastest light response, we generated transgenic flies expressing this version of NLS-LARIAT for *in vivo* testing.

**Figure 1.**
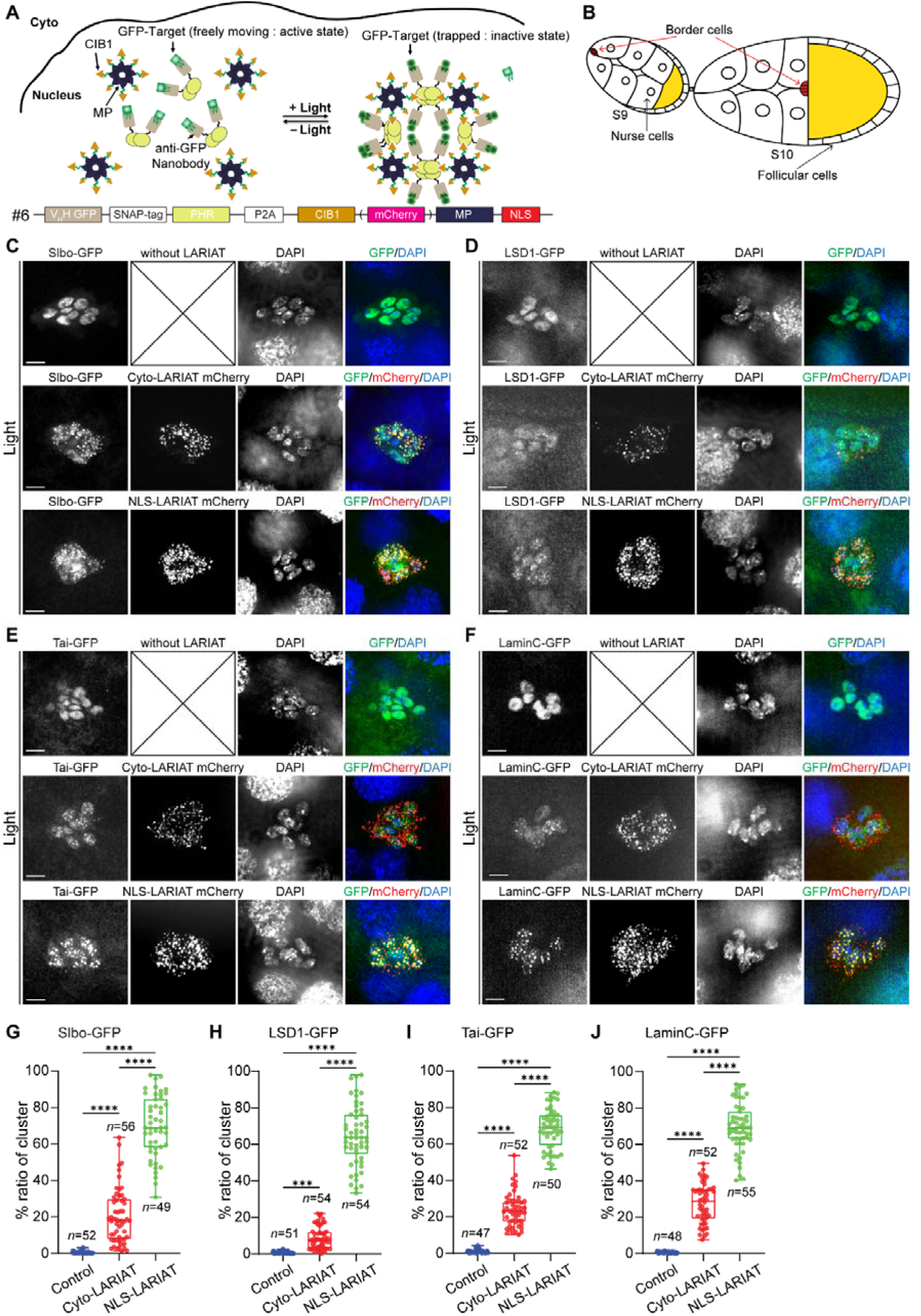
NLS-LARIAT strongly clusters various GFP-tagged nuclear factors in border cells. **(A)** Schematic summarizing the working mechanism of NLS-LARIAT. The construct shown here corresponds to version #6, which was used in subsequent *Drosophila* experiments. **(B)** Diagram of *Drosophila* stage 9 (S9) and 10 (S10) egg chambers showing follicle cells, border cells, and nurse cells. **(C** to **F)** Representative images of border cells expressing Slbo-GFP (**C**), LSD1-GFP (**D**), Taiman-GFP (**E**) and LaminC-GFP (**F**) without LARIAT, with Cyto-LARIAT-mCherry or with NLS-LARIAT-mCherry, under the light illumination. DNA was stained with DAPI. **(G** to **J)** Quantifications of the clustering percentage (clustered GFP signals relative to total GFP signals) in border cells with Slbo-GFP (**G**), LSD1-GFP (**H**), Taiman-GFP (**I**), and LaminC-GFP (**J**), with the indicated background under the light illumination. *n* represents the total number of border cell groups examined. ***p < 0.001; ****p < 0.0001; one-way ANOVA with Kruskal–Wallis test followed by Dunn’s multiple comparisons test. Scale bars are 10LJμm in (**C** to **F**). Boxplots show 25th and 75th percentiles (boxes), minimum and maximum values (whiskers), and all data points (dots) from biologically independent samples in (**G** to **J**).

Next, we used *Drosophila* ovary egg chamber as an *in vivo* system to test this new optogenetic tool. *Drosophila* ovary contains 15 strings of egg chambers at different developing stages from stage 1 to stage 14 (*25*). The egg chamber is composed of a monolayer follicular epithelium surrounding a 16-cell germline cyst (Fig. 1B; (*25*)). *Drosophila* follicle epithelial cells have been used to study the controlling mechanisms of cell division and tumor growth (*26, 27*). The tight coordination between follicle cells and germline cyst has been used to study intercellular communication within a tissue (*28*). From early stage 9, a migratory group of epithelial cells detaches from the epithelia and basement membrane, migrates through germline nurse cell junctions, and reaches the oocyte (Fig. 1B). This process, named border cell migration, has been used to study epithelial-mesenchymal transition and invasive collective cell migration (*25, 29, 30*). Our previous studies have demonstrated that border cells and follicle cells can be used to study spatiotemporal roles of factors and signals in controlling collective cell migration and actomyosin pulsatile contractility, respectively (*18, 19, 31, 32*).

Here, we compared the clustering efficiency of NLS-LARIAT vs. Cyto-LARIAT (by overnight light illumination) for different GFP-tagged nuclear factors in *Drosophila* border cells, follicle cells and nurse cells by using Slbo-Gal4, Tj-Gal4, and Nanos-Gal4 to drive UAS-LARIAT. These GFP-tagged nuclear factors include transcription factors (Slbo [*slow border cells*] and Taiman), an epigenetic factor (histone demethylase LSD1 [Lysine-specific demethylase-1]) and nuclear envelope proteins (Lamin C and Kugelkern). Slbo is highly expressed in border cells, Taiman is highly expressed in border cells and follicle cells, and LSD1, Lamin C and Kugelkern are expressed in all three cell types. We used the percentage ratio of cluster (see Methods) to quantify the clustering efficiency for this comparison. Compared with the control, light-stimulated Cyto-LARIAT had little-to-mild clustering effects on most GFP-tagged nuclear factors in border cells and follicle cells (Fig. 1, C to J; and fig. S3, A to E). In contrast, light-stimulated NLS-LARIAT had very robust clustering effects on all these nuclear factors in all three cell types (Fig. 1, C to J; and fig. S3). These results confirm that NLS-LARIAT is suitable for clustering-trapping various GFP-tagged nuclear factors in the *Drosophila in vivo* system.

### NLS-LARIAT phenocopies genetic inhibition of border cell migration

We next asked whether NLS-LARIAT−mediated clustering-trapping of GFP-targets could produce a photo-inhibitory effect. To perform this test, we applied NLS-LARIAT to border cell migration because it is a powerful system that allows to study efficient and rapid optogenetic manipulation of various cytoplasmic factors (*31, 32*). We characterized the photo-inhibitory effects of NLS-LARIAT on a GFP-tagged transcriptional factor (Slbo-GFP) or an epigenetic factor (LSD1-GFP) during border cell migration and compared them with the inhibition of either factor by RNAi. Slbo is a transcription factor known to determine border cell migratory fate; genetic loss-of-function mutation of *slbo* strongly blocks the formation of border cell group, the detachment of border cells from follicular epithelium, as well as the migratory ability of border cells (*33*). LSD1 is the first identified histone demethylase that demethylates histone H3K4 (*34*); genetic loss-of-function mutation of *lsd1* leads to *Drosophila* ovarian defects (*35, 36*). However, whether LSD1 controls border cell migration has not been explored yet. Here, genetic knockdown of Slbo and LSD1 by expressing their RNAi in border cells resulted in strong migration defects (showing different levels in migration delay calculated by the migration index) as well as some detachment defects, compared with the control (fig. S4, A to H); and live cell imaging also showed that the migratory speed of these two RNAi-expressing border cells was significantly reduced (fig. S4, I to L).

We then compared the global clustering-trap effects of NLS-LARIAT on either Slbo-GFP or LSD1-GFP during border cell migration. To achieve a global cluster-trap effect, we kept flies periodically illuminated for at least 6 hours (to accommodate the entire period of border cell migration) and also during tissue dissection to allow the clustering process to persist before the experiment (see Methods). Fixed imaging showed that compared with border cells not expressing LARIAT, light-stimulated NLS-LARIAT caused significant defects in the migration and detachment abilities of either Slbo-GFP or LSD1-GFP expressing border cells (Fig. 2, A to D), which is similar to the migration and detachment defects observed in either RNAi-expressing border cell groups. However, NLS-LARIAT in the dark did not cause border cell migration defects (Fig. 2, A to D). In contrast to severe migratory defects induced by NLS-LARIAT clustering-trap, light-stimulated Cyto-LARIAT had no significant impact on the migratory ability of border cells expressing Slbo-GFP or LSD1-GFP: almost all these Cyto-LARIAT-expressing border cell groups arrived at the oocyte during stage 10, as calculated by the migration index (fig. S4, M to P). Thus, these results support that light-stimulated clustering-traps of critical transcriptional factors and epigenetic factors by NLS-LARIAT, but not by Cyto-LARIAT, can phenocopy the effects of genetic inhibition on border cell migration. Next, we performed live cell imaging to obtain the dynamic migration behavior induced by NLS-LARIAT−mediated photo-inhibition (the clustering process occurred before the experiment as fixed imaging analysis). Live imaging results demonstrated that, compared with the control, border cells with light-stimulated NLS-LARIAT−mediated Slbo-GFP clustering (Movie, S1 and 2) or LSD1-GFP clustering (Movie, S3 and 4) exhibited a much slower migration speed (Fig. 2, E to H), similar to the dynamic migration defects observed in border cells expressing either RNAi. These live cell imaging results further confirm that the NLS-LARIAT−mediated photo-inhibition of key nuclear factors resembles genetic inhibition of border cell migration.

**Figure 2.**
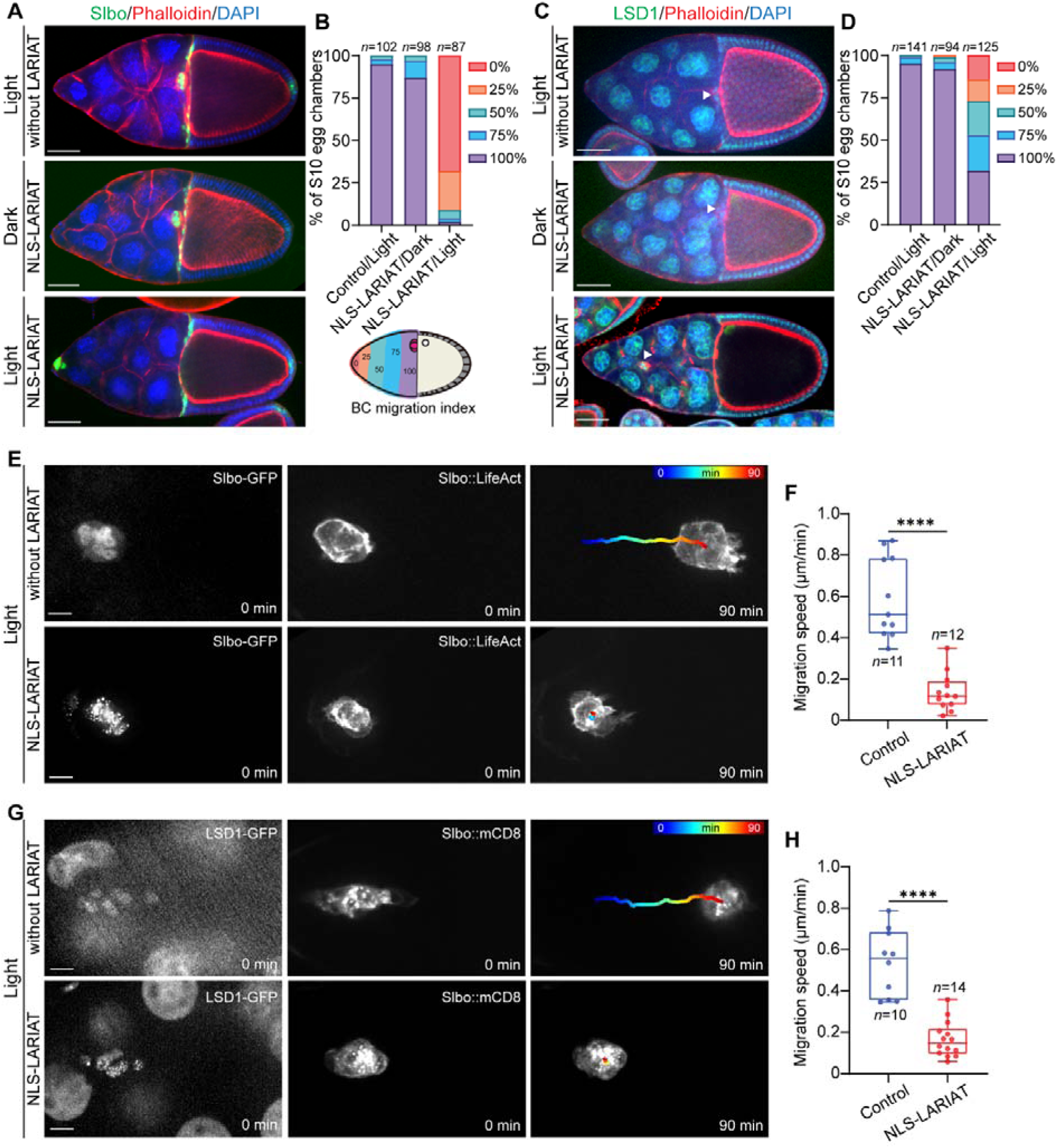
Clustering-trap of Slbo-GFP and LSD1-GFP inhibits border cell migration. **(A** and **C)** Representative images of Slbo-GFP−expressing **(A)** and LSD1-GFP−expressing **(C)** egg chambers at stage 10. Upper panels: without NLS-LARIAT in border cells under the light condition; middle and lower panels: with NLS-LARIAT expression in border cells under dark and light conditions, respectively. F-actin and nuclei were visualized with phalloidin and DAPI staining. In **(C)**, the arrowhead indicates the border cell cluster, as LSD1-GFP signal is relatively weak in border cells. **(B** and **D)** Quantifications of border cell migration index (summarized in cartoon) in Slbo-GFP–expressing (**B**) and LSD1-GFP–expressing (**D**) egg chambers at stage 10. Stacked bar plots show the percentage of egg chambers exhibiting 0%, 25%, 50%, 75%, or 100% completion of border cell migration. *n* represents the total number of egg chambers examined. **(E** and **G)** Representative time-lapse images of Slbo-GFP expressing (**E**) and LSD1-GFP expressing (**G**) border cell groups without (upper) or with (lower) NLS-LARIAT under the light. Slbo::LifeAct-RFP and Slbo::mCD8-RFP were used to monitor F-actin and membrane signals in migrating border cells. Time points shown are 0 and 90 min; colored tracks indicate migration trajectories. **(F** and **H)** Quantifications of the migration speed of Slbo-GFP–expressing (**F**) and LSD1-GFP–expressing (**H**) border cell groups with or without NLS-LARIAT under the light. *n* represents the total number of border cell groups examined. ****p < 0.0001, nonparametric Mann–Whitney U test. Scale bars are 50LJμm in (**A** and **C**) and 10LJμm in (**E** and **G**). 25th and 75th percentiles as box limits, minimum and maximum values as whiskers; each datapoint is displayed as a dot (from n biologically independent samples) in (**F** and **H**).

Given the similar inhibitory phenotypes, we asked whether light-stimulated clustering-traps might similarly affect downstream targets of Slbo or LSD1. Here, we tested downstream targets for which commercially available antibodies from the Developmental Studies Hybridoma Bank (DHSB) are available. Previous studies reported that Slbo can positively affect the transcription of Singed (Sn, encoding a member of the fascin family of actin-binding proteins), Tenascin major (Ten-m, encoding a transmembrane protein), Upheld (Up, encoding the striated muscle protein Troponin T), Actin(57B) (*37, 38*). Indeed, we found that Slbo-GFP clustering border cells exhibited a drastic reduction in protein levels of all these genes, compared with the control border cells (Fig. 3, A to H). Our previous transcriptomic studies, performed in ovaries where LSD1 was depleted by RNAi, showed that LSD1 can negatively regulate the transcription of Sidestep (side, encoding a transmembrane protein of the immunoglobulin superfamily) and Ten-m (*39*). Here, we found that LSD1-GFP clustering border cells exhibited significantly increased levels of these two proteins (Fig. 3, I to L). These results thus confirm that photo-inhibition of Slbo-GFP or LSD1-GFP by clustering-traps similarly affects the expression levels of Slbo or LSD1 downstream targets in border cells, as genetic inhibition does. Altogether, our findings support that light-stimulated NLS-LARIAT can effectively phenocopy genetic manipulations, offering a robust approach for studying gene function in border cell migration. Interestingly, Slbo-GFP clusters were prominently exported to the cytoplasm, while LSD1-GFP clusters remained confined to the nucleus (fig. S5, A and B). As Slbo-GFP signals clustered under periodical blue light illumination, small clusters always appeared within the border cell nucleus, then many clusters were located along the nuclear envelopes, and finally large clusters were enriched in the border cell cytoplasm (fig. S5, C and D; and Movie S5). It thus supports the idea that the clustering process mediated by light-stimulated NLS-LARIAT might accelerate the shuttling of Slbo-GFP from the nucleus to the cytoplasm, suggesting a role for intrinsic nuclear localization and export signals in regulating the dynamic behavior of aggregated proteins.

**Figure 3.**
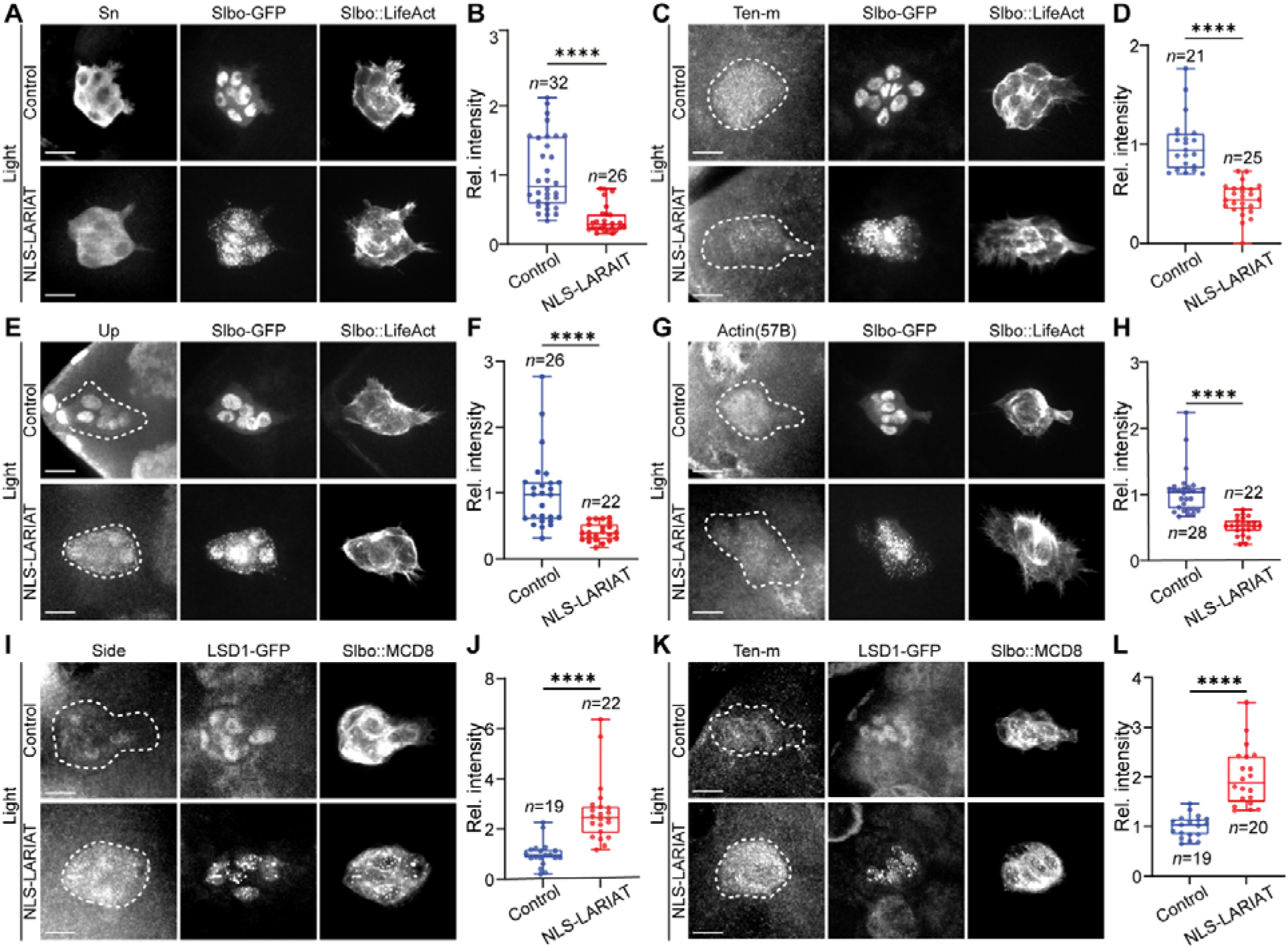
Confirmation of the effects of Slbo-GFP and LSD1-GFP clustering on target gene expression in border cells. **(A, C, E, G, I** and **K)** Representative images of Sn (**A**), Ten-m (**C**), Up (**E**) and Actin(57B) (**G**) in Slbo-GFP expressing border cells, and Side (**I**) and Ten-m (**K**) in LSD1-GFP expressing border cells, without or with NLS-LARIAT co-expression under the light. Sn, Ten-m, Up, Actin(57B), and Side were detected by antibody staining. Slbo::LifeAct-RFP and Slbo::MCD8-RFP were used to monitor the migratory behaviors of Slbo-GFP expressing and LSD1-GFP expressing border cells. **(B, D, F, H, J** and **L)** Quantifications of relative signal intensity of Sn (**B**), Ten-m (**D**), Up (**F**) and Actin(57B) (**H**) in Slbo-GFP expressing border cells, of Side (**J**) and Ten-m (**L**) in LSD1-GFP expressing border cells, without or with NLS-LARIAT, under the light. Control values were normalized to 1. *n* represents the total border cell groups examined. ****p < 0.0001, the nonparametric Mann–Whitney U test (B, D, F, H and L) and two-tailed Student’s t test (J). Scale bars are 10LJμm in (**A, C, E, G, I** and **K**). Boxplots show medians, 25th and 75th percentiles (boxes), minimum and maximum values (whiskers), and all data points (dots) from biologically independent samples in (**B, D, F, H, J** and **L**).

In addition to our study of border cell migration, we also investigated the long-term photo-inhibitory effect of NLS-LARIAT on LSD1-GFP in follicle cells during egg chamber development, since LSD1 is known to control the normal development of egg chambers (*35*). Here, continuous periodic blue light illumination was applied 6 to 24 hours to allow the clustering process to occur prior to the assay (see Methods). Light illumination led to strong LSD1-GFP clusters in follicle cells expressing NLS-LARIAT, compared with normal LSD1-GFP distribution in control cells (fig. S6A). Consequently, NLS-LARIAT expressing females laid significantly fewer eggs compared to controls after 24-hour photoactivation (fig. S6, B and C). Moreover, the developmental competence of the eggs was severely impaired. While eggs in the control group efficiently developed into larvae, the hatching rate of the larvae in the NLS-LARIAT group began to decline sharply as early as 6 hours after light exposure and gradually decreased after 12 hours and 24 hours of light illumination (fig. S6, B and D). These phenotypes strongly resemble *lsd1* loss-of-function mutants and demonstrate that sustained sequestration of nuclear LSD1 impairs both egg production and developmental potential. Together, these results establish NLS-LARIAT as a versatile tool for long-term photo-inhibition of nuclear factors to explore their roles in cell fate decision and tissue development.

### NLS-LARIAT reveals spatiotemporal control of border cell migration

Optogenetic tools allow us to understand more rapid effects of target proteins compared to the long-term effects of genetic manipulations. Transcriptional activation has previously been regarded as a long-time control, while recent studies imply that it can be achieved within a short time and a few optogenetic tools (e.g. iLexy) have been established to rapidly control gene expression (*14*). We first determined the temporal role of Slbo-GFP clustering-trap by gradual photo-inhibition. *Drosophila* was cultured, and then tissues were dissected and mounted in the dark, so that Slbo-GFP clustering would not occur at the beginning of live cell imaging. Then, to better observe the Slbo-GFP clustering process and its mediated gradual changes in border cells, we photo-treated the entire border cell group every 3 minutes (see Methods). Upon repeated blue light illumination, Slbo-GFP clustering progressively increased, accompanied by a gradual reduction in border cell migration speed, while the control border cells without Slbo-GFP clustering showed normal migration speed (Fig. 4, A to D; and Movie, S6 and 7). The migration speed of border cells seemed to respond to the clustering process within 15-30 minutes (Fig. 4D), indicating that Slbo clustering-trap can rapidly control border cell migration ability. As Slbo-GFP clusters increased, we observed the gradual appearance of large and excessive protrusions (Fig. 4, B and E), and notably these protrusions were frequently observed in border cells expressing Slbo RNAi as well as expressing Slbo-iLEXYi under the dark condition (fig. S7, A and C). It thus suggests that temporal photo-inhibition of Slbo-GFP can influence border cell protrusions to disturb border cell migration ability. Different from NLS-LARIAT, three versions of LEXY (Slbo-LEXY, Slbo-iLEXYi and Slbo-iLEXYs, with different recovery times) all exhibited very robust nuclear-to-cytosol shuttling process even in darkness, inhibiting the action of Slbo in border cells (fig. S7, A and B). It thus implies that the Slbo-LEXY tool has a strong leaking effect in the dark state, and that NLS-LARIAT is more suitable than iLEXY for temporally dissecting certain nuclear factors with acute nuclear export signals. In addition, temporal Slbo-GFP clustering led to severe defects in the border cell detachment process (fig. S8), consistent with previous observations of *slbo* loss-of-function mutants (*33*). All these results support that light-stimulated NLS-LARIAT can rapidly induce the clustering-trap and photo-inhibition of key transcriptional factors (e.g. Slbo), thereby resulting in corresponding changes in migratory border cells.

**Figure 4.**
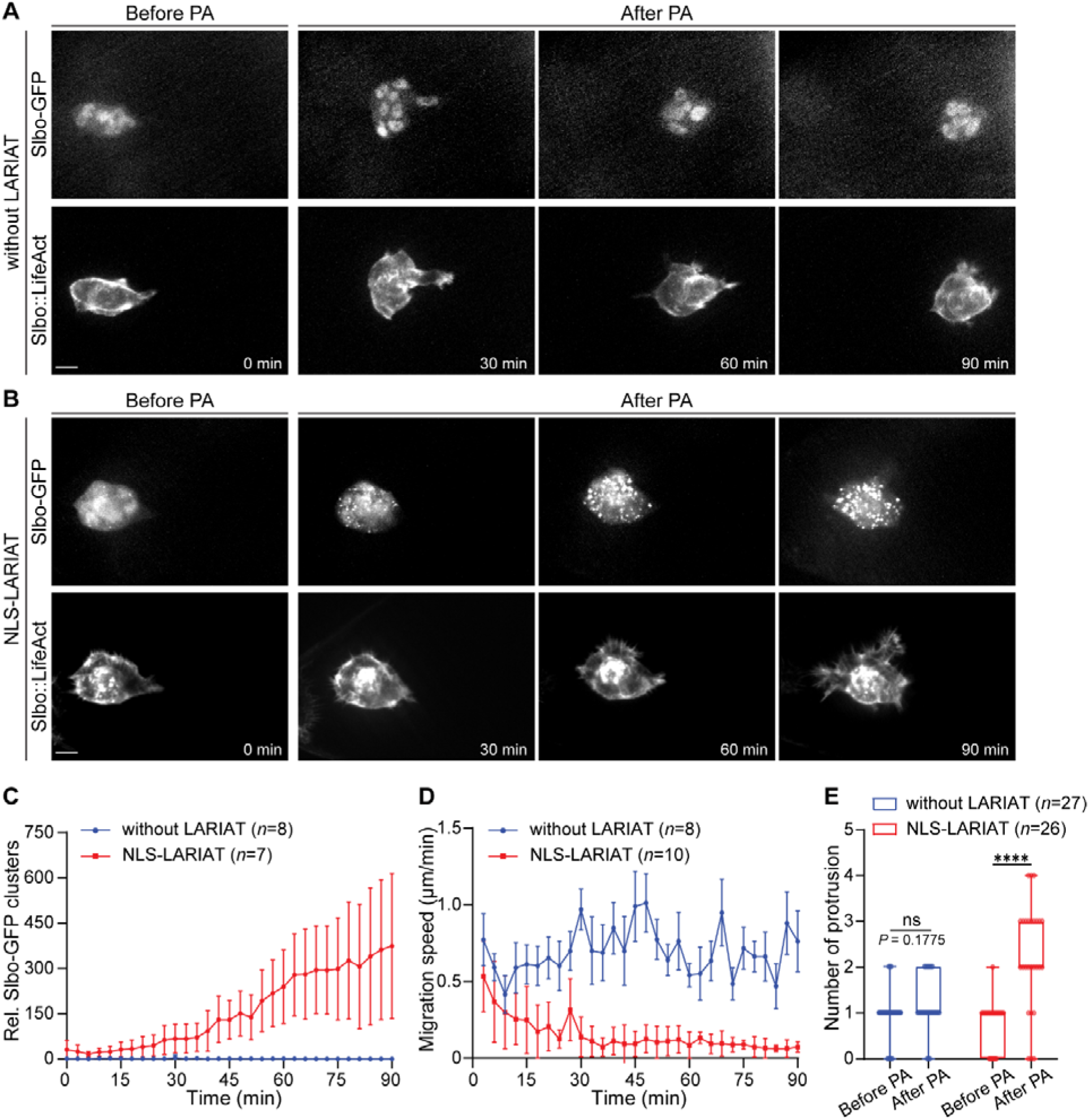
Gradual clustering of Slbo-GFP rapidly blocks border cell migration. **(A** and **B)** Representative time-lapse images of Slbo-GFP expressing border cell groups without (**A**) or with (**B**) NLS-LARIAT, before and after blue light illumination. Slbo::LifeAct-RFP was used to monitor F-actin signals in migrating border cells. **(C)** Time-lapse quantification of Slbo-GFP clusters in Slbo-GFP expressing border cell groups, without NLS-LARIAT or with NLS-LARIAT, before and after blue light illumination. **(D)** Time-lapse quantification of mean migration speed (μm per minute) in Slbo-GFP expressing border cell groups, without NLS-LARIAT or with NLS-LARIAT, before and after blue light illumination. **(E)** Quantification of protrusion numbers in Slbo-GFP expressing border cell groups without NLS-LARIAT or with NLS-LARIAT before and after blue light illumination. ****p < 0.0001; ns, not significant, nonparametric Mann–Whitney U test. Scar bar is 10 μm in (**A** and **B**). Data in (**C** and **D**) are presented as mean ± SD. *n* represents the sample size. Boxplots show medians, 25th and 75th percentiles (boxes), minimum and maximum values (whiskers), and all data points (dots) in (**E**).

Next, we characterized the spatial effects of NLS-LARIAT on localized photo-inhibition of Slbo-GFP in the migrating border cell group. In contrast to temporal experiments, here we photo-treated rear or leader border cells every 10 seconds to test their rapid responses to localized NLS-LARIAT−mediated photo-inhibition. Local activation of NLS-LARIAT in rear border cells of the Slbo-GFP expressing group rapidly induced the protrusion growth at the photo-treated cells, while leader border cells gradually lost their original leading protrusions (Fig. 5, A and B), indicating that ectopic protrusion growth in rear cells can antagonize protrusions in other border cells; differently, this localized photo-treatment had no effect on rear cells of the control border cell group (Fig. 5, A and B). This result indicates that strong Slbo levels in border cells might locally inhibit protrusion growth. To further verify this conclusion, we photo-activated NLS-LARIAT in leader cells without prominent leading protrusions, and we observed that it rapidly induced prominent protrusions in photo-treated leader cells, whereas local photo-treatment had no effect on the control leader cells without protrusions (Fig. 5, C and D). Consistent with these local photo-inhibitory effects, in wild-type border cell groups, leader border cells with large protrusions exhibited significantly lower levels of nuclear Slbo-GFP than follower border cells during migration; interestingly, some leader border cells gradually reduced their nuclear Slbo-GFP signals as new protrusions grew out (Fig. 5, E and F; Movie S8). Slbo is considered as a core factor controlling border cell fate and migration ability, but its precise role in migrating border cell groups after detachment remains unclear. Unexpectedly, our localized photo-inhibitory effects contradict the logical speculation that nuclear Slbo promotes border cell migration through its ability to control efficient protrusion growth, but support the hypothesis that differential levels of nuclear Slbo in border cells (moderate levels in leaders vs. high levels in followers) are crucial for the distribution of polarized protrusion in the border cell group, enabling efficient and directional collective movement.

**Figure 5.**
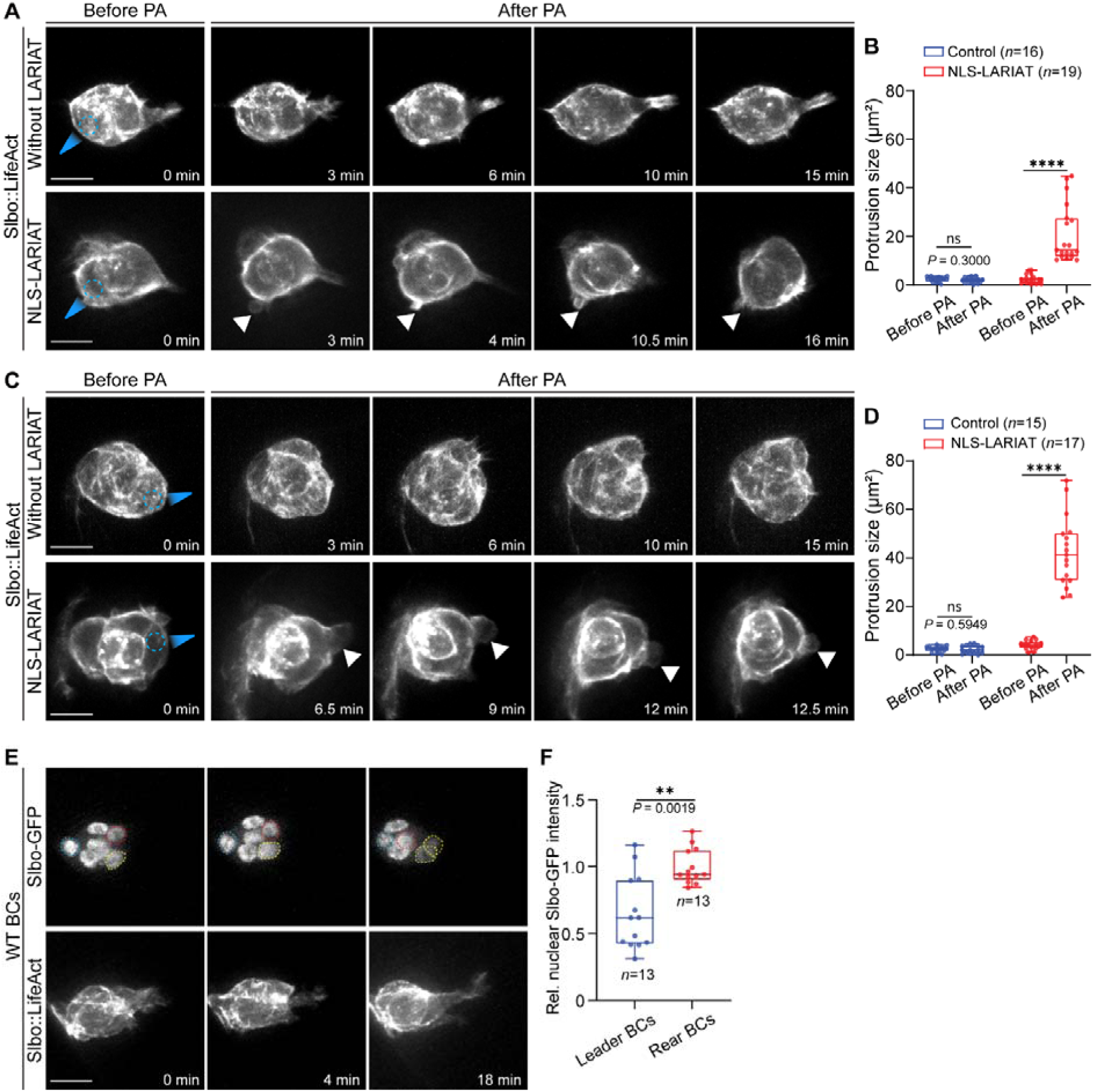
Local photo-inhibition of Slbo-GFP reveals an inhibitory role for Slbo in border cell protrusions. **(A** and **C)** Representative time-lapse images of Slbo-GFP expressing border cell groups, without (upper panels) or with (lower panels) NLS-LARIAT, before and after local blue light illumination at rear border cells (**A**) and leader border cells (**C**). Slbo::LifeAct-RFP was used to monitor F-actin signals in migrating border cells. Blue arrowheads mark blue light illumination region and white arrowheads indicate protrusions. **(B and D)** Quantifications of protrusion size in rear (**B**) and leader (**D**) indicated border cells before and after local photo-treatment. *n* represents the total sample size. Statistical significance was assessed by the nonparametric Mann–Whitney U test. ****p < 0.0001; ns, not significant. **(E)** Representative time-lapse images of Slbo-GFP and Slbo::LifeAct-RFP in the border cell group with large leading protrusions. Dotted blue, red, and yellow circles mark rear border cells, leader border cells with large protrusions at the beginning, and leader border cells without protrusions at the beginning, respectively. **(F)** Quantification of relative nuclear Slbo-GFP intensity in leader cells with large protrusions, compared with rear border cells without protrusions. The average value in rear border cells was normalized to 1. *n* represents the total sample size. Statistical significance was assessed by the nonparametric Mann–Whitney U test. **p = 0.0019. Scar bar is 10μm in (**A, C** and **E**). Boxplots show medians, 25th and 75th percentiles (boxes), minimum and maximum values (whiskers), and all data points (dots) from biologically independent samples in (**B, D** and **F**).

Taken together, our results highlight the advantage of NLS-LARIAT for precise, temporal, and local control of gene function during border cell migration.

### NLS-LARIAT modulates nuclear envelope and chromatin structure

Almost all available optogenetic tools are limited to transcription factors and their activities, with few reports on their application in controlling other nuclear proteins/structures (*5*). Nuclear structures, such as the nuclear envelope and chromatin structure, play important roles in protecting DNA and maintaining genome integrity. The former is mechanically supported by networks of lamin filaments (e.g. Lamin B) (*40*); these lamin proteins are essential components of the nuclear lamina, providing structural support and regulating nuclear shape and stiffness (*41, 42*). The latter is supported by histone proteins, highly conserved proteins that play critical roles in chromatin structure and function; histone proteins associate with DNA to form nucleosomes and contribute to the regulation of transcription and DNA replication (*43*). Thus, we asked whether light-stimulated NLS-LARIAT can photo-manipulate structural nuclear proteins, such as nuclear envelopes (lamin B) or histone proteins by clustering-trapping.

In follicle cells expressing only Lamin B-GFP, the nuclear envelope typically exhibited a smooth contour with a natural curvature along the rim (Fig. 6A). In contrast, in follicle cells co-expressing Lamin B-GFP and NLS-LARIAT, photo-activation induced the progressive aggregation of Lamin B-GFP along the nuclear rim. After several hours of periodic blue light illumination (3–6 hours), nuclear envelopes gradually became more rounded and irregular, and following prolonged light treatment (9-14 hours), obvious bubble-like protrusions appeared (Fig. 6A). Super-resolved images from random illumination microscopy (RIM) confirmed that these morphological abnormalities corresponded to nanoscale Lamin B clusters embedded in the nuclear envelope (Fig. 6B). To investigate whether Lamin B clustering alters nuclear membrane mechanics, we employed optical tweezers to directly measure nuclear stiffness in Lamin B-GFP normal vs. clustering follicle cells (Fig. 6C). NLS-LARIAT–induced clustering significantly increased nuclear membrane stiffness, reflected by elevated restoring forces compared with the control (Fig. 6D). Notably, most Lamin B-GFP clusters remained stably associated with the nuclear rim and were unable to dissociate (Fig. 6, A and B), likely condensing lamin filaments locally and thereby reinforcing the nuclear lamina. Together, these results demonstrate that NLS-LARIAT enables spatiotemporal photo-manipulation of nuclear lamina architecture and mechanics, providing an optogenetic strategy to dissect the role of lamins in nuclear membrane organization and mechanics *in vivo*.

**Figure 6.**
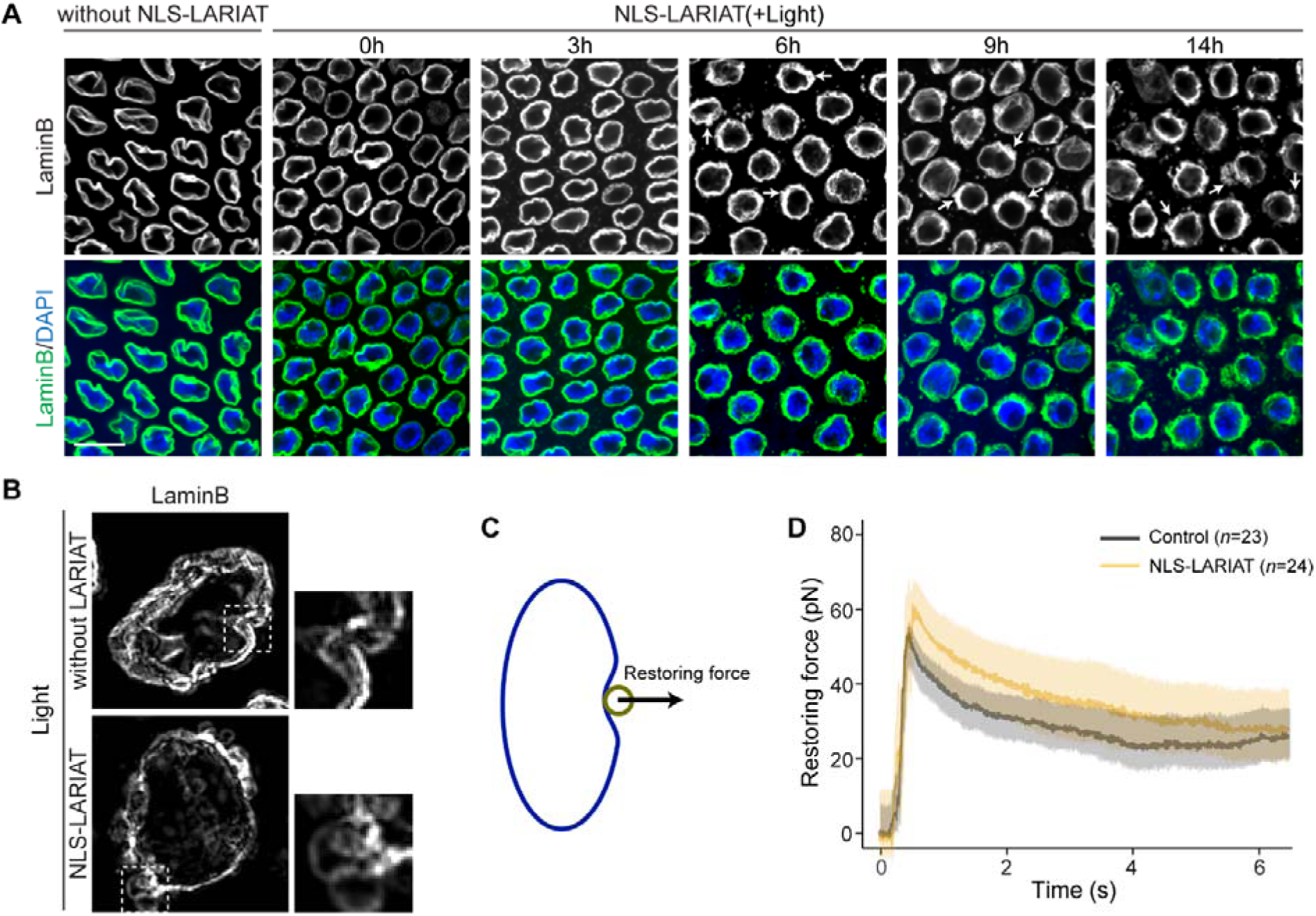
Lamin B-GFP clustering modulates nuclear membrane stiffness in epithelial cells. **(B)** Representative images of LaminB in follicle cells without NLS-LARIAT (left) and with NLS-LARIAT under blue light illumination at the indicated time points (0, 3, 6, 9, and 14 hours). Lamin B-GFP labelled nuclear membrane of follicular cells was shown in green and nuclei were labeled with DAPI (blue). Arrows at different time points indicate disrupted nuclear envelope morphology. Scale bar, 10 μm. **(C)** Representative RIM super-resolution images of Lamin B-GFP expressing follicle cells, without (upper panels) and with (lower panels) NLS-LARIAT, under blue light illumination (14 hours). Dotted squares mark the regions that are magnified in the right panels in (**B**). **(D)** Schematic illustrating the measurement of restoring force by optical tweezers. A calibrated displacement was applied to the cell membrane, and the restoring force was recorded. **(E)** Time course of the restoring force measured by optical tweezers in follicular cells under blue light illumination, comparing Control (n=23, gray) and NLS-LARIAT-expressing cells (n=24, yellow). Cells were subjected to a calibrated displacement, and the restoring force was recorded over time. Shaded areas represent mean ± SEM.

Given that follicle cells require 1 or 2 days to change their fate, to allow clustering to occur, we subjected flies harbouring His2A-GFP in all cell types and NLS-LARIAT expression in follicle cells to periodic blue light illumination for 28 hours prior to the experiment. Photo-activation induced robust His2A-GFP clustering that was colocalized with mCherry-tagged NLS-LARIAT and DNA, indicating that clustering did not disrupt the native histone–DNA association (fig. S9A). Strikingly, in egg chambers containing His2A-GFP clusters under the light state (but not in tissues under the dark state), follicle cells exhibited less cell number and bigger nuclear size, whereas germline nurse cells showed more cell number (more than 15 nurse cell per egg chamber) and smaller nuclear size (fig. S9). In contrast, under the same periodic light illumination, control egg chambers expressing either His2A-GFP alone or NLS-LARIAT alone developed normally, with typical cell number and normal nuclear size (fig. S9). These findings suggest that sustained histone clustering disrupts egg chamber homeostasis by either blocking mitosis or triggering a premature initiation of the endocycle in follicle cells, and driving aberrant germline proliferation and/or disrupting germline/follicle cells communication and eggs chamber encapsulation.

## Discussion

Optogenetic approaches for nuclear factors are limited to transcription factors and are rarely applicable to other nuclear factors and structures (*5*). Here, we established a novel optogenetic tool named NLS-LARIAT, which can be broadly applied to virtually every nuclear factor in *Drosophila* to study nuclear processes in *in vivo* system. NLS-LARIAT can cluster different GFP-tagged nuclear proteins (e.g. transcription factors, epigenetic factors, nuclear membrane proteins, etc.) in *Drosophila* ovarian follicle cells, border cells and nurse cells. NLS-LARIAT−mediated clustering-trap of Slbo or LSD1 phenocopies the effects of genetic inhibition on migrating border cells, while it can reveal gradual, rapid, and localized inhibitory effects on border cell migration, thus outperforming genetic manipulation. Importantly, NLS-LARIAT is also suitable for the perturbation of other nuclear factors, such as nuclear membrane components and histones.

Unlike currently available optogenetic approaches for nuclear factors (*5*), NLS-LARIAT is based on the clustering-trap strategy that can recognize GFP targets localized in the nucleus. Thus, NLS-LARIAT can be applied to any GFP target, allowing the application of a common optogenetic tool to different nuclear factors, eliminating the need to redesign optogenetic tools when the target changes. This is the first advantage of NLS-LARIAT compared to other optogenetic tools for nuclear proteins. This clustering-trap strategy also enables the optogenetic control of stable nuclear factors and their associated structures to study various nuclear processes *in vivo* beyond transcription/epigenetics, which is impossible with other optogenetic approaches. Our successful photo-manipulation of LaminB and Histone2A corroborates this second advantage of NLS-LARIAT. Compared with other optogenetic approaches (*12, 14*), NLS-LARIAT showed slow and partial recovery in the dark after the clustering process, in contrast to the faster reversibility observed with Cyto-LARIAT (*20*). We suspect that this reflects the crowded, constrained nuclear environment (limited free volume, chromatin anchoring, higher viscoelasticity) that impedes cluster dispersion. Thus, our tool might not be suitable for experiments requiring rapid, repeated activation–recovery cycles, but it matched up well with acute loss-of-function and sustained sequestration *in vivo*.

Remarkably, clustering-traps of different nuclear factors in *Drosophila* egg chambers exhibit distinct phenotypes. 1) Clustering-trap of the transcriptional factor (Slbo-GFP) and epigenetic factor (LSD1-GFP) phenotypically mimics the effects of their genetic inhibition on border cell migration, including similar migration defects and similar changes in the expression of their downstream targets. Indeed, Slbo-GFP clusters are gradually exported to the cytosol, supporting the photo-inhibition of this transcription factor via dissociating the Slbo protein from its functional sites (genomic regions where Slbo binds). 2) Differently, clusters of Lamin B (a key component of nuclear membrane structure) cannot dissociate from the nuclear envelope, and thus this clustering-trapping causes the nuclear membrane to stiffen rather than soften. We speculate that Lamin B clustering might locally increase the concentration of the protein, thereby stiffening the nuclear envelope. Our tool thus provides an optogenetic strategy for spatiotemporally regulating nuclear membrane stiffness, which might also be applicable to nuclear envelope modulation in other models. Moreover, Histone2A clusters might not dissociate from DNA and chromosomes, but these clusters seem to disturb cell fate in some way: perhaps by affecting 3D chromatin structure thus influencing various processes from transcription to DNA replication and perturbing cell proliferation (*44–46*). Here, the distinct phenotypes of clustering-trapping different nuclear factors appear to be highly dependent on the dynamics and nano-structures of GFP-targets: more dynamic proteins are more prone to dissociate from their functional location by clustering-trap and completely lose the function, whereas more stable proteins could remain in their original location and structure even after clustering-trapping, leading to other perturbations rather than loss of function. For more stable proteins (e.g. Lamin B and Histones), new optogenetic strategies will need to be designed to dissociate these proteins from their functional sites.

Our studies demonstrate that NLS-LARIAT can be used for short-term or long-term photo-manipulation of GFP-targets. Interestingly, clustering-trap of Slbo-GFP can rapidly induce local protrusion changes in migrating border cells, while inhibiting protrusions in other border cells. In contrast, long-term clustering-trap of Slbo-GFP results in the appearance of ectopic large protrusions in all border cells, hampering the migratory ability of border cells. The difference between short-term and long-term photo-inhibition suggests that partial loss of Slbo-GFP might produce a phenotype that is distinct from Slbo strong clustering-trapping and complete loss of Slbo function. Thus, it will be interesting and important to compare the photo-inhibitory effects of short-term and long-term clustering-traps of different GFP-tagged nuclear factors in border cells and follicle cells, as well as in any type of cells and tissues in other models.

In summary, NLS-LARIAT provides a new optogenetic strategy for the photo-manipulation of a variety of GFP-tagged nuclear factors. It can be applied to photo-manipulate various nuclear factors and structures, over varying time periods to produce short- or long-term effects, and to locally photo-disturb diverse targets within a migratory group or a tissue. Although NLS-LARIAT has not yet been applied to mammalian cultured cells and other *in vivo* models (e.g. mouse, zebrafish, Xenopus, etc.), we anticipate that adapting it to other expression vectors will readily enable photo-manipulation of GFP-tagged nuclear factors, thereby facilitating precise spatiotemporal studies of their roles across diverse models and systems.

## Materials and Methods

All materials and methods are available in the online supplementary information.

## Acknowledgments

We thank Bloomington and VDRC *Drosophila* stock center for flies. We thank Karine Belguise for the discussion of manuscript preparation. This work is supported by Agence Nationale de la Recherche (ANR PRC AAPG2022 NEW-corset), Scientifiques de la Fondation ARC (PJA20191209714) and 2020 du Canceropole GSO to X.W.; Agence Nationale de la Recherche (ANR-20-CE12-0015-01, COMETES) to L.D.S.; National Natural Science Foundation of China (8240082473) to S.Z.; J.L. is supported by ANR PRC AAPG2022 NEW-corset. B.L. was supported by a PhD fellowship from the China Scholarship Council (CSC) (Grant No. 202204910032) and Fondation ARC pour la Recherche sur le Cancer (Grant No. ARCDOC42025010009296).

## Author contributions

B.L., L.D.S. and X.W. designed the project. B.L., S.Z., H.L., and Y.G. performed experiments. A.G. generated Slbo::LEXY, Slbo::GFP and NLS-LARIAT transgenic flies. D.J. generated LSD1::GFP transgenic flies. B.L. and J.L. developed methods and data analysis scripts and analysed the data. S.Z. and T. M. performed optical tweezers. T.M. analysed the restoring force from optical tweezers. H.L. performed RIM super-resolution microscopy. B.L., L.D.S. and X.W. wrote the manuscript.

